# Adaptation to an invasive host is collapsing a native ecotype

**DOI:** 10.1101/030858

**Authors:** M L Cenzer

## Abstract

Locally adapted populations are often used as model systems for the early stages of ecological speciation, but most of these young divergent lineages will never become complete species. While the collapse of incipient species is theoretically common, very few examples have been documented in nature. Here I show that soapberry bugs (*Jadera haematoloma*) have lost adaptations to their native host plant (*Cardiospermum corindum*) and are regionally specializing on an invasive host plant (*Koelreuteria elegans*), collapsing a classic and well-documented example of local adaptation. All populations that were adapted to the native host - including those still found on that host today - are now better adapted to the invasive in multiple phenotypes. Weak differentiation remains in two traits, suggesting that homogenization across the region is incomplete. This study highlights the potential for adaptation to invasive species to disrupt native communities by swamping adaptation to native conditions through maladaptive gene flow.

## Main Text

Locally adapted populations are often used as models for the early stages of speciation, particularly in the recent discussion of speciation driven by differential ecological conditions^1^. These lineages are convenient for the study of divergent evolution because they can arise quickly, sometimes in tens or hundreds of generations, and the drivers of differentiation are often contemporary and identifiable. However, they are also theoretically more susceptible to collapse than young species due to incomplete intrinsic reproductive isolation^2^. The maintenance of local adaptation often relies on spatial isolation or continuing selection against migrants and hybrids, processes that are highly dependent on the stability of the local environment. Changes in the quality^3^, abundance, stability^4,5^, and proximity of habitats can all theoretically drive hybridization and collapse of young divergent lineages.

Existing examples of young species collapse fall into two broad categories: (1) If lineages have been diverging in allopatry and come into secondary contact before speciation is complete, one lineage may be absorbed into the other. In at least two cases, endemic species have been lost via this process^6^. (2) Environmental disturbance may break down reproductive barriers between lineages already co-existing in sympatry and produce a hybrid swarm^7-9^. Two examples of this process are the collapse of over 100 species of cichlids in Lake Victoria^8^ and the more local collapse of a benthic-limnetic stickleback species pair^9^. This mechanism is also responsible for the breakdown of beak size divergence in one population of Galapagos finches^10^. Cases where collapse occurs *prior* to the completion of speciation, for example, during local adaptation, are exceptionally rare.

I analyzed populations of soapberry bugs that were locally adapted to different host plants in 1988 to determine whether these populations are following an ecological speciation trajectory or collapsing back together.

The red-shouldered soapberry bug *Jadera haematoloma* (Hemiptera: Rhopalidae) has provided a textbook case of local adaptation. Soapberry bugs are native to the southern peninsula and Keys of Florida, where they have co-evolved with a native balloon vine (*Cardiospermum corindum*). In the mid-1900s, the golden rain tree (*Koelreuteria elegans*) was widely introduced to the peninsula of Florida and colonized by soapberry bugs^11^. By 1988 there was clear evidence of host-associated local adaptation. Adult feeding morphology was locally adapted to differences in host seedpod size: adult females adapted to *C. corindum* had long beaks hypothesized to be for feeding through the large seedpods of that host, while females from populations on *K. elegans* had short beaks suitable for feeding through the flattened pods of this host^11^ (Fig. 1a). Juveniles had high survival and short development time on their local host and reduced survival and prolonged development time on the alternative host^12,13^ (Fig. 2a & Fig. 3a). Females from *C. corindum* produced relatively few large eggs and females from *K. elegans* produced many small eggs^12^ (Fig. 4a). Based on this evidence of strong local adaptation and the largely allopatric distribution of the two host plants in Florida, this system was a strong candidate for ecologically driven ‘incipient speciation’.

**Figure 1.**
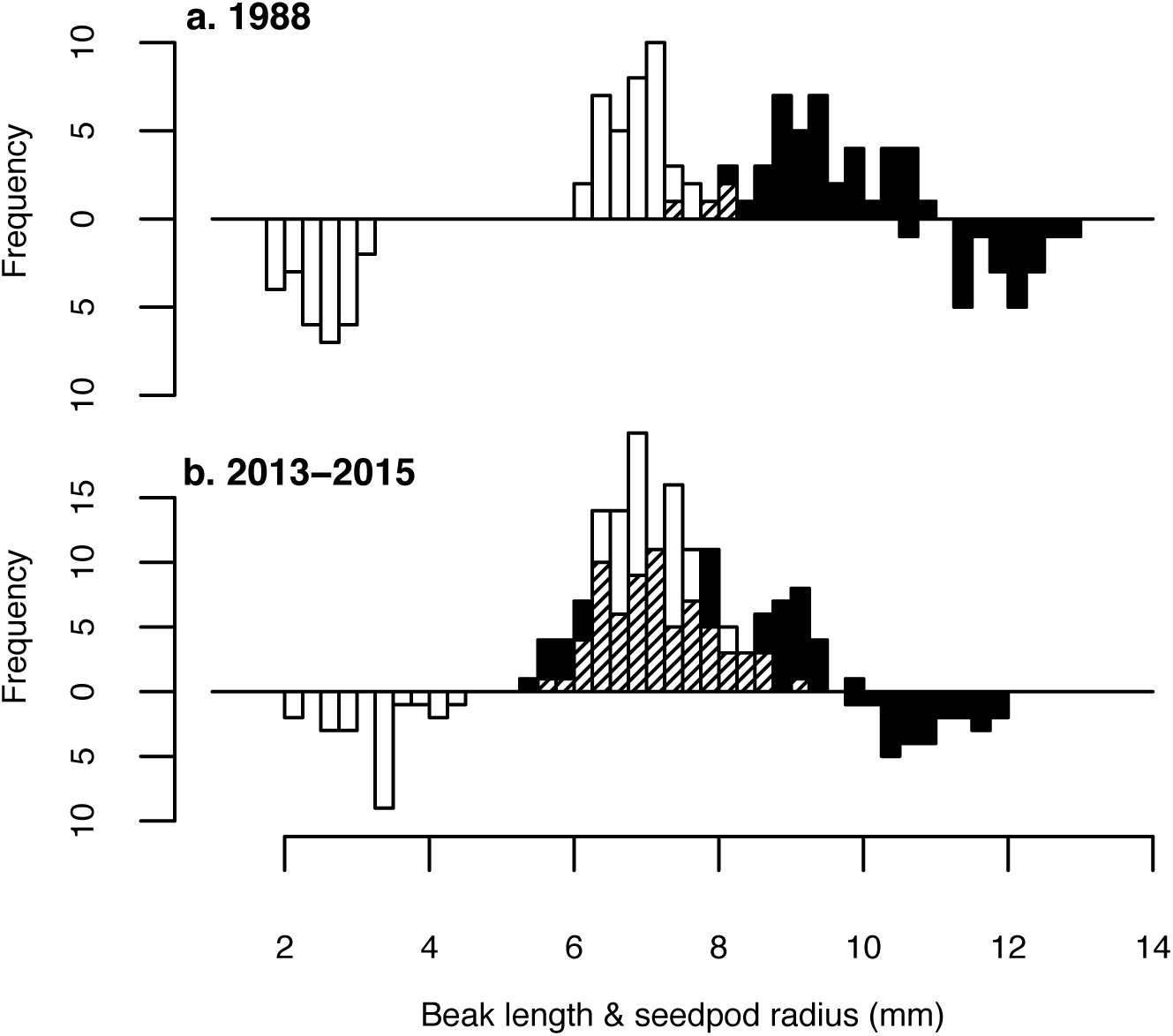
Beak length and pod size on both hosts between years. Black bars indicate pods or bugs collected from *C. corindum,* white bars indicate pods or bugs from *K. elegans,* and cross-hatching indicates areas where the two distributions are overlapping. a. 1988 female *J. haematoloma* beak length (upright bars) and pod size of each host (inverted bars). Data taken with permission from Carroll & Boyd 1992. b. Combined 2013-2015 female *J. haematoloma* beak length (upright bars) and pod size of both host plants (inverted bars). Data for 7 additional sites collected in 2013-2015 is available in Extended Data Fig. 1.

**Figure 2.**
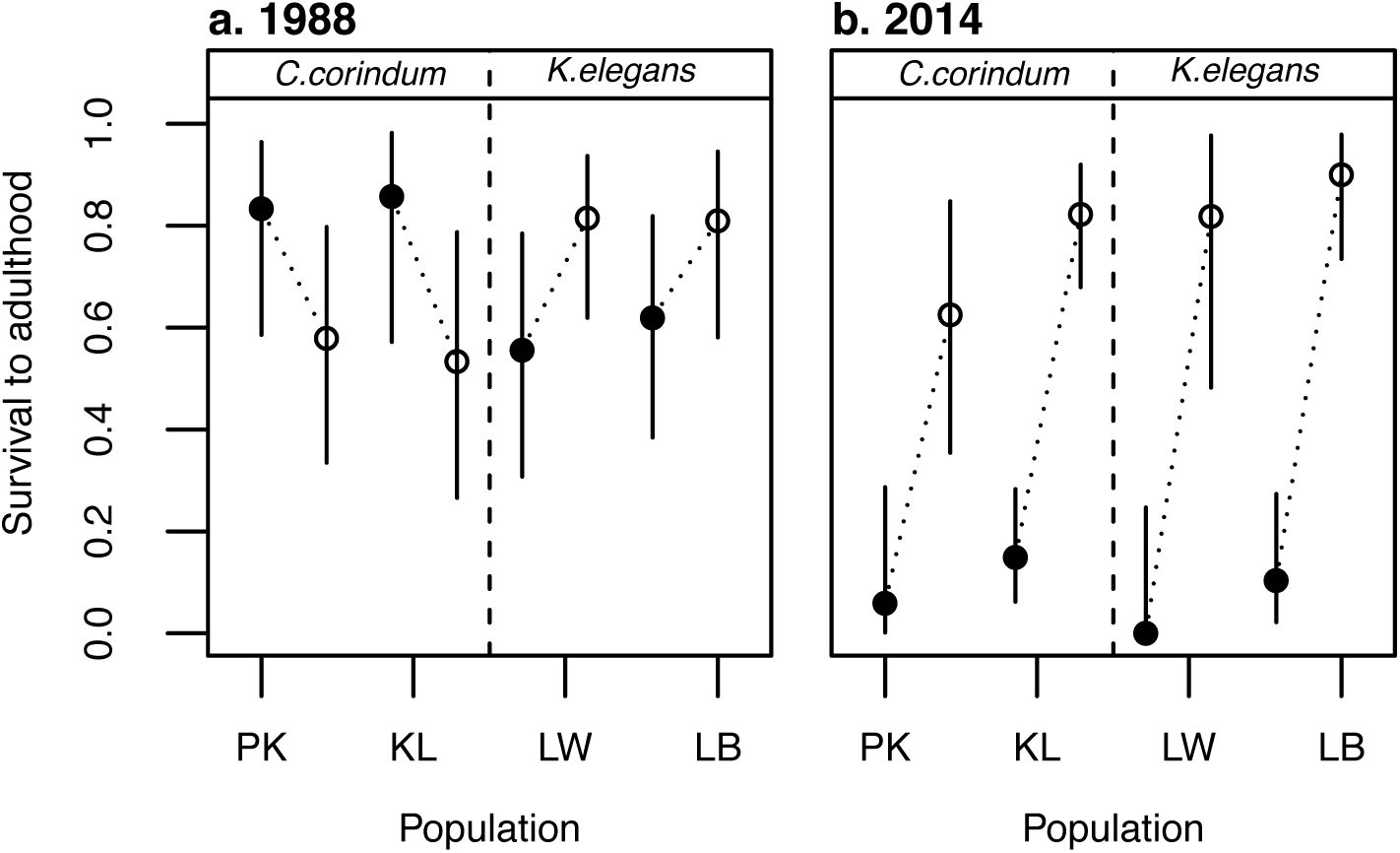
Juvenile survival on both host plants in 1988 and 2014. a. Proportion of juveniles from 4 populations surviving to adulthood when reared on *C. corindum* (black circles) and *K. elegans* (white circles) in 1988. Data taken with permission from Carroll et al 1998. b. Proportion of juveniles from the same 4 populations surviving to adulthood when reared on both host plants in 2014. Populations PK and KL were collected from *C. corindum;* LW and LB from *K.elegans.* Error bars represent the 95% binomial confidence interval using the Pearson-Klopper method. Data for 4 additional sites in 2014 is available in Extended Data Fig. 2.

**Figure 3.**
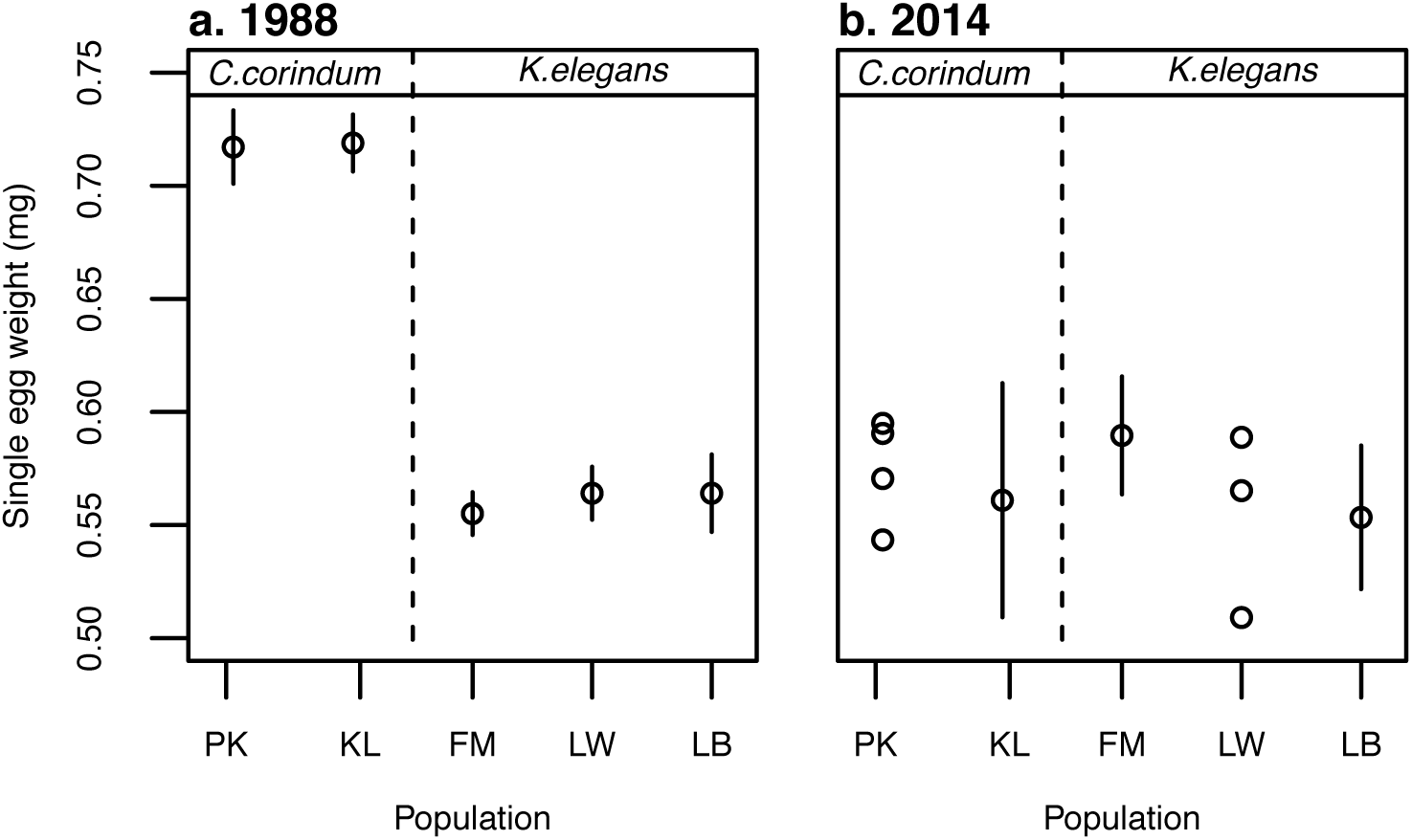
Individual egg weight differentiation in 1988 and 2014. a. The average weight of individual eggs laid by females from 5 different populations in 1988. Data taken with permission from Carroll et al 1998. b. The average weight of individual eggs laid by females from the same 5 populations in 2014. Populations PK and KL were collected from *C. corindum;* FM, LW and LB from *K.elegans.* Due to limited sample size, all points for PK and LW are plotted individually. Error bars represent 95% confidence intervals. Data for 3 additional sites in 2014 is available in Extended Data Fig. 3.

**Figure 4.**
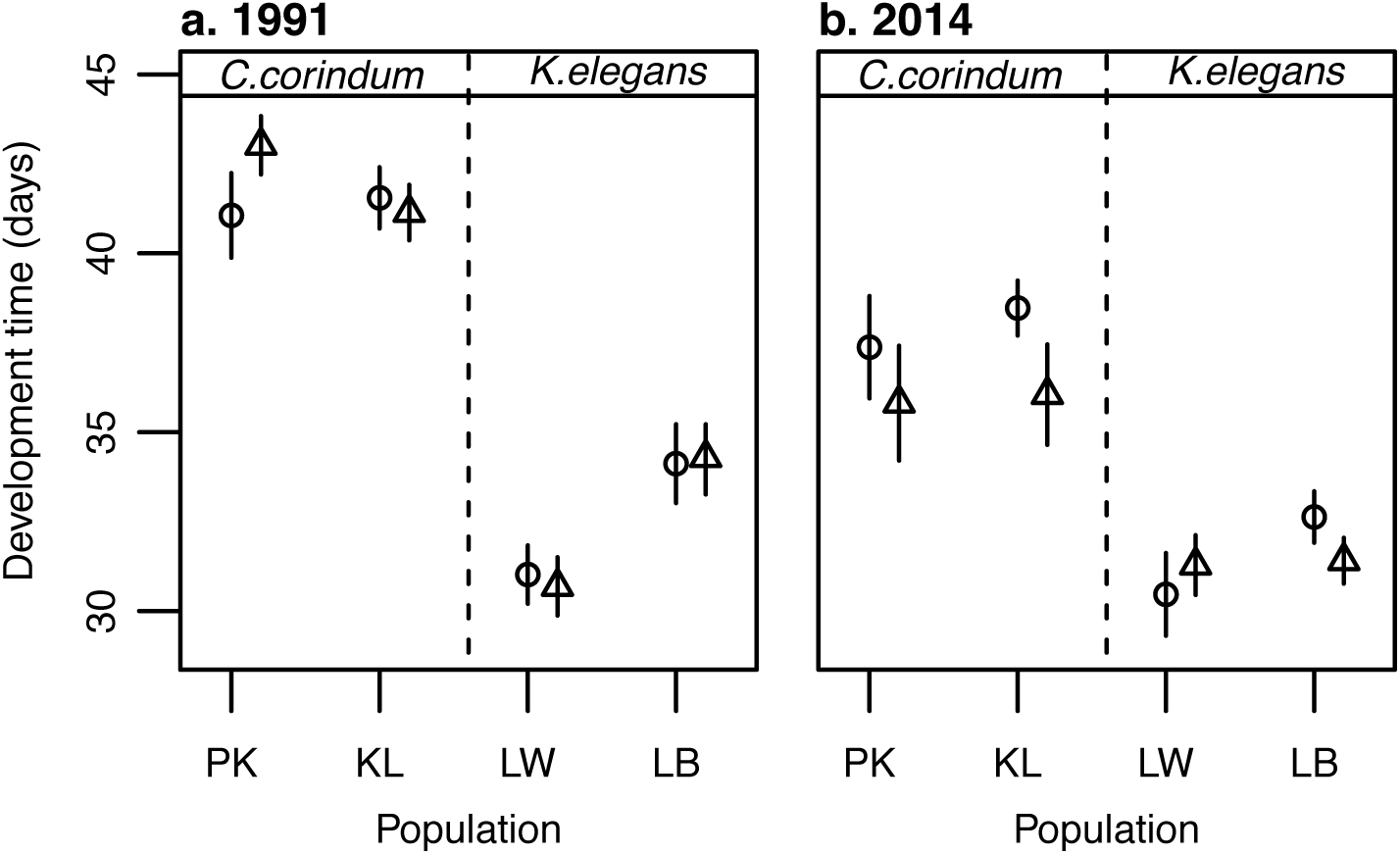
Development time in 1991 and 2013-2014. a. Development time for females (circles) and males (triangles) from 4 populations in 1991. Data taken with permission from Carroll et al 1997. b. Development time for the same 4 populations combined from 3 experiments in 2013 and 2014. Populations PK and KL were collected from *C. corindum;* LW and LB from *K.elegans.* All individuals were reared on *K.elegans.* Error bars represent 95% confidence intervals. Data for 4 additional sites in 2013-2014 and by experiment is available in Extended Data Fig. 4. Data for individuals reared on *C.corindum* is presented in Extended Data Fig. 5.

## Results

Current evidence demonstrates that soapberry bugs in Florida are not following a speciation trajectory, but have instead become more similar in all traits that were previously locally adapted. In 1988, the mean beak length of bugs on the native *C. corindum* was 34% greater than that of bugs on the invasive *K. elegans* (9.33±0.21mm vs. 6.93±0.12mm; t-value=16.02, p=<.0001, df=68). In 2013-2015, the difference between the same two host-associated populations was reduced to 5% (7.48±0.18mm vs. 7.13±0.10mm; t-value=2.724, p=.007, df=170) (Fig. 1). In a total of 8 populations across Florida measured in 2013-2015, the difference in beak length between the two hosts was not statistically significant (7.32±0.12mm vs. 7.15±0.06mm; t-value=-1.57, p>.05, df=514; Extended Data Fig. 1). This reduction in differentiation is driven by changes on the native host plant. Beak lengths of individuals in Key Largo, where *C. corindum* associated beak length was originally estimated, have decreased to 80% of their historical length (t-value=10.978, p<.0001, df=104). In contrast, bugs collected in Lake Wales, where *K. elegans-associated* beak length was originally estimated, have shown very little change since 1988 (t-value=-1.952, p=.054, df=97). The observed beak length change is in the opposite direction predicted by the hypothesis that beak length should be adapting to match pod size in these populations.

Cross-rearing experiments also support the hypothesis that contemporary populations are less differentiated than they were historically (Fig. 2). In 2014, survival from hatching to adulthood was dramatically higher on the invasive host *K. elegans* than on *C. corindum* for all populations (0.81 vs. 0.10; z-value=8.908, p<.0001; Fig. 2b). This is in stark contrast to historical cross-rearing experiments, in which juvenile survival was consistently higher on the host from which a population was collected (interaction z-value=3.299, p<.001; Fig. 2a).

Changes in individual egg weight also support a loss of local differentiation (Fig. 3). The mean egg weight for two populations from *C. corindum* was 0.712±0.018mg per egg in 1988, while populations on *K. elegans* had eggs averaging 0.560±0.012mg per egg. In 2014, egg weight averaged 0.565±0.036mg per egg in the same two *C. corindum* populations and 0.570±0.019mg per egg in the same three *K. elegans* populations (t-value=.205, p=.84, df=19).

This reduction in differentiation is driven entirely by changes in the egg weight of individuals from *C. corindum* (t>5, df=18, p<.001), while egg weight for individuals from *K. elegans* has remained unchanged (t<1.76, df=48, p>0.05). This lack of differentiation is consistent across a total of 8 populations measured in 2014 (0.573±0.023 [*C.corindum*] vs. 0.572±0.015 [*K.elegans*]; t-value=0.076, p=0.94, df=36; Extended Data Fig. 3).

Changes in development time are also consistent with decreased differentiation (Fig. 4). In 1991, individuals from *C. corindum*-associated populations had a mean development time of 41.69±0.94 days while those from *K. elegans*-associated populations had a mean development time of 32.53±0.78 days when both were reared on *K. elegans* (t-value=-13.762, p<.001). In 2013 and 2014, the same populations from *C. corindum* took an average of 37.21±0.62 days to reach adulthood, while those from *K. elegans* took 31.48±0.33 days (t-value=-11.511, p<.001) when reared on *K. elegans.* There has been a pronounced decrease in development time in *C. corindum*-associated populations (year effect=4.61 days; t-value=7.37, p<0.001, df=149) that is absent in *K. elegans*-associated populations (year effect=.86 days; t-value=1.64, mean p=.2, df=206). The pattern of reduced, but detectable, differentiation is consistent across the 8 total populations measured in 2013-2014 (35.29±0.49 days vs. 31.58±0.27 days; t-value=10.8, p<.001, df=246; Extended Data Fig. 4).

## Discussion

This classic case of local adaptation is transitioning to a set of populations all specialized on the invasive *K. elegans.* The simplest explanation for this pattern is that gene flow from *K. elegans* to *C. corindum* has increased over the last 26 years. This case does not conform to either of the existing mechanisms of species collapse, both of which require sympatry. Instead, genes facilitating survival on *K. elegans* may have followed a ‘stepping stone’ process of migration from one *C. corindum* population to the next as a response to increased migration from *K. elegans.* This was likely facilitated by the increasing abundance of the invasive host on the lower peninsula and decreasing regional abundance of the native host^11^. Stronger competition on *C. corindum* from native herbivores that are not contending with maladaptive gene flow may be exacerbating the negative effects of decreased local adaptation on abundance.

Although there are few empirical examples, theory predicts the degradation of local adaptation under a wide range of environmental conditions^12,13^. In 2001, Ronce and Kirkpatrick predicted a downward spiral of maladaptation and population size, termed ‘migrational meltdown’, occurring in spite of strong local selection under moderate migration, resulting in a single specialist if migration is asymmetric^14^. This theory suggests that maladapted populations of soapberry bugs on *C. corindum* are unlikely to be ‘rescued’ by evolution^15^, but will likely remain sinks unless migration from *K. elegans* decreases well below earlier rates.

The current literature on local adaptation is heavily skewed toward cases that appear to be progressing towards speciation, potentially because these are the cases that persist long enough to be detected. As environmental disturbance rapidly shifts the targets of selection, local adaptation is likely to emerge and collapse in many systems as populations move entirely onto novel resources.

The rapid adaptation of soapberry bugs to *K. elegans* in 1988 contradicted the general expectations of evolutionary ecology as one of the seminal examples of evolution occurring over ecologically relevant time scales^16^. The second evolutionary shift in this system, the extensive loss of adaptation to their ancestral host plant throughout the state of Florida, is similarly striking. Adaptation on short time scales has proven to be more of a rule than an exception^17-21^. The ephemerality of local adaptation may prove to be similarly common.

## Methods

### Collection

*Jadera haematoloma* were collected in May 2013, December 2013, April 2014, and April 2015 from the following locations (coordinates of sample collection sites in parentheses; directions available on request): Gainesville (26.660477, -82.346965), Leesburg (28.796398,-81.877980), Lake Wales (27.934719, -81576640), Ft. Myers (26.633952, -81.879978), Homestead (25.570843, -80.455236 [*C.corindum*] & 25.551041, -80.420876 [*K.elegans*]), Key Largo (25.175560, -80.367695), and Plantation Key (24.991368, -80.539964). The northern four sites were collected from *Koelreuteria elegans* and the southern two sites were collected from *Cardiospermum corindum*; bugs were collected from both host plants in Homestead, the only presently known sympatric site. Host plant seeds were collected from each site in December 2013 and April 2014 and stored at 4°C until they were used for rearing. Seeds with indications of previous feeding were discarded. Each *K. elegans* site had 5-10 individual trees and each *C. corindum* site had 3-15 individual vines sampled.

### Morphology

Adults from the field were phenotyped to the nearest .02 mm for beak length (the distance from the anterior tip of the tylus to the distal tip of the mouthparts), thorax width (taken at the widest part of the pronotum), and forewing length (from anterior to distal tip) using Mitutoyo calipers under a dissecting microscope. A subset of adults had the additional measurement of body length, the length from the anterior tip of the tylus to the distal tip of the closed wings. Female morphology is reported in Fig. 1 and Extended Data Fig. 1, Fig. 6 & Fig. 7. Since differentiation in any male morphology has not been previously documented, and was not found here, these results are not discussed. Male morphological data will be publicly archived along with female data.

### Sample sizes

For 1988 data, sample size for female beak length on *C. corindum* and *K. elegans* was 44 and 40, respectively. Sample size for pod size on *C. corindum* and *K. elegans* was 20 and 28, respectively. For 2013-2015 data, sample size for female beak length on historically measured *C. corindum* and *K. elegans* was 107 and 109, and sample size for pod radius on *C. corindum* and *K. elegans* was 19 and 22, respectively. Sample sizes for additional populations (Plantation Key, Homestead (*C. corindum*), Homestead (*K. elegans*), Ft. Myers, Leesburg, and Gainesville) were 34, 43, 29, 88, 82, and 32, respectively. Sample sizes were maximized to the extent that availability in the field and time permitted, with additional weight being given to populations with available historical data. All individuals that were collected were measured unless they were so physically damaged in transport as to make morphology unreliable (<2% of all individuals collected). No blinding was done during these measurements.

### Cross-rearing

All rearing was carried out in Sanyo Versatile Environmental Test Chambers at 28°C during the day and 27.5°C at night, 50% relative humidity with a 14:10 Light:Dark cycle, following climate conditions from Carroll et al 1998. Adults from the field were housed in vented Petri dishes lined with filter paper and given water in a microcentrifuge tube stoppered with cotton (“water pick”) and 3 seeds of their field host plant. Eggs were collected daily until hatching; adults were then frozen at -20C for morphological analyses. Nymphs were removed within 12 hours of hatching to reduce egg cannibalism and housed individually in mesh-lidded cups lined with filter paper with a water pick and a seed of their randomly assigned host plant. Four nymphs were used from each mother; two were assigned to each host plant. Prior to hatching, datasheets were generated that randomized the order of the location of origin for each seed using the “sample” command in R. When the first nymph in a family hatched, a coin was flipped to determine which host it would be assigned to; the second hatching nymph was assigned to the opposite host. The same procedure was carried out for the third and fourth nymphs. This was to ensure that there would not be a bias in early hatching nymphs being assigned more frequently to one host or the other, and to avoid having more individuals reared on one host or the other in the event that families had fewer than 4 eggs successfully hatch. Additional seeds (a total of 2 for *K.elegans* and 3 for *C.corindum,* for a total seed mass of ~150mg) were added at 7 days of age. Individual containers were rotated daily within mesh boxes (each holding 36 individuals), and boxes were rotated daily within the growth chamber to reduce the effect of specific location. Water, paper and cotton were changed weekly. Nymph survival was assessed daily.

### Sample sizes

For 1988 data, sample sizes for cross-rearing in each population (Plantation Key, Key Largo, Lake Wales, and Leesburg) are n=14, 18, 18, and 21 reared on *C. corindum,* and n=15, 19, 27, and 21 reared on *K. elegans.* For 2014 data, sample sizes for the same populations are n=17, 47, 13, and 29 reared on *C. corindum* and n=16, 45, 11, and 30 reared on *K. elegans.* Each population was represented by n=8, 23, 8, and 15 independent families. For the 4 populations not represented in historical datasets (Homestead (*C. corindum*), Homestead (*K.elegans*), Ft. Myers, and Gainesville) the sample sizes are n=34, 7, 36, 36, and n=34, 7, 35, and 39 reared on *C. corindum* and *K. elegans* respectively. Each of these populations is represented by n=17, 4, 18, and 21 independent families. Sample sizes were limited by 1) the availability of adults at each field site; 2) survival of collected adults through transport within Florida and from Florida to the lab where rearing was conducted; and 3) whether or not individual females produced viable eggs after transport. Females were housed only with seeds from their local site both during transport and during egg production; therefore, any selection that may have occurred during this process should have exacerbated local adaptation rather than masking it. No blinding of observers was used in this metric.

### Egg mass

Individual egg mass was collected from surviving females reared in the F1 laboratory generation (rearing methods described above). Females were randomly paired with males from the same population, but with different parents, within 2-5 days of eclosion. Individuals were not paired earlier to allow time for the exoskeleton to harden, reducing susceptibility to cannibalism and physical damage to the genitalia during mating attempts. Water, paper, and cotton were changed weekly, and two additional seeds were added weekly. Eggs were counted and massed in groups (from 1 to 84 eggs per group) to within .01 mg daily for the first 10 days after a female began laying eggs, resulting in up to 10 separate estimates of egg mass per female. Individuals who did not lay eggs in the first 30 days after reaching adulthood were excluded, as that is the estimated life expectancy in the field (Carroll 1991). The egg mass reported is combined for individuals reared on both *K. elegans* and *C. corindum.*

### Sample sizes

For 1988 data, sample sizes for each population (Plantation Key, Key Largo, Ft. Myers, Lake Wales, and Leesburg) are n=16, 20, 24, 16, and 50. For 2014 data, sample sizes for the same 5 populations are n=4, 9, 13, 3, and 12. For the 3 populations without historical data (Homestead (*C. corindum*), Homestead (*K. elegans*), and Gainesville), sample sizes are n= 8, 2, and 13. Sample sizes for this metric were primarily restricted by mortality during development, the limited number of potential mates within the appropriate population and time frame, and willingness of individuals to mate and produce eggs. All individuals that successfully reproduced were included in this metric; however, females who eclosed later were slightly less likely to be included, because in some cases all available males had already been assigned to earlier eclosing females by the time they emerged. This problem was in part due to the fact that males had slightly shorter development time than females in this experiment, a pattern that was not present in previous studies, and was therefore not anticipated during mate allocation. No blinding of observers was used in this metric.

### Development time

Development time was combined from three separate experiments using the rearing methodology described above. The first experiment was conducted in May 2013 (Extended Data Fig. 4b) the second from the F1 generation in April 2014 (Extended Data Fig. 4c), and the third from the F2 generation in the same year (Extended Data Fig. 4d). Individuals were checked daily for eclosion to adulthood, and every individual that survived was included. Development time is only reported in the main text for individuals reared on golden rain tree. Very few individuals successfully reached adulthood on balloon vine in each of these experiments, so there is potential for a single generation of strong selection driving observed patterns on this natal host. Mortality occurred largely during the first instar. This is likely due to either small beak size or weak beak musculature in first instar bugs, which become less problematic once bugs reach later instars. It is possible that individuals with short first instars were therefore more likely to survive, which could contribute to a shortening of overall development time in individuals reared on balloon vine. Consistent with this hypothesis, the development times of individuals surviving to adulthood on balloon vine were significantly shorter than those developing on golden rain tree (Extended Data Fig. 5b); however, given the scarcity of these data, I do not draw any conclusions here. No blinding of observers was used in this metric.

### Sample sizes

For 1991 data, sample sizes for each population (Plantation Key, Key Largo, Lake Wales, and Leesburg) were n=15, 24, 35, and 33, for females and n=15, 26, 42, and 43 for males. For the same populations in 2013-2014, sample sizes are n=16, 34, 15, 27, for females and n=16, 20, 21, and 27 for males. These metrics were combined from 3 experiments, each of which followed the same protocol, but had slightly different sample sizes for each population. In 2013, data was collected for 7 populations (Plantation Key, Key Largo, Homestead (*C. corindum*), Homestead (*K. elegans*), Ft. Myers, Lake Wales, and Leesburg) with n=7, 4, 9, 8, 5, 11, and 6 for females and n=12, 4, 15, 10, 10, 11, and 2 for males. In 2014, generation 1, data was collected for 8 populations (Plantation Key, Key Largo, Homestead (*C. corindum*), Homestead (*K. elegans*), Ft. Myers, Lake Wales, Leesburg, and Gainesville) with n=8, 23, 12, 2, 16, 4, 11, and 14 for females and n=2, 14, 20, 3, 16, 7, 17, and 15 for males. In 2014, generation 2, data was collected for 8 populations (Plantation Key, Key Largo, Homestead (*C. corindum*), Homestead (*K. elegans*), Ft. Myers, Lake Wales, Leesburg, and Gainesville) with n=1, 7, 5, 2, 11, 0, 10, and 13 for females and n=2, 2, 7, 1, 11, 3, 8, and 11 for males. For individuals reared on *C.corindum* in 1991, sample size for each population (Plantation Key, Key Largo, Lake Wales, and Leesburg) were n=15, 24, 35, and 33 for females and n=15, 26, 42, and 43 for males. For all experiments combined in 2013-2014, sample sizes for each population (Plantation Key, Key Largo, Homestead (*C. corindum*), Homestead (*K. elegans*), Ft. Myers, Lake Wales, Leesburg, and Gainesville) is n=11, 7, 9, 3, 2, 0, 4, and 4 for females and n=3, 4, 5, 2, 3, 4, 4, and 1 for males. Sample size was limited by the same factors that limited collection for assessing survival, and then by mortality itself, as development time can only be measured for individuals that reach adulthood.

### Statistical analyses

All analyses were conducted in the statistical program R version 3.2.2. Analyses including random factors used the package lme4. Testing assumptions for homoscedasticity used the package lmtest. Binomial confidence intervals were generated using the package binom.

### Beak length analyses

Historical individual beak length measures and corresponding body sizes were located in the field notebooks of Scott Carroll from the year 1988, when they were originally collected.

Linear models (*lm*) with all combinations of host, year, body size, and all possible interactions (18 possible models) with Gaussian error distributions were compared using AIC. For this analysis, the years 2013, 2014 and 2015 were grouped and compared to 1988 as discrete factors, rather than as a continuous variable. To assess how year and host influence beak length individually, pairwise comparisons were conducted using Welch’s two-sample t-test, the results of which are reported in the main text. Beak length in 2014 fails to reject normality using the Shapiro-Wilk normality test; in 1988, beak length on each host plant individually fails to reject normality using the Shapiro-Wilk normality test.

### Correcting for body size

Body size was controlled for in Carroll & Boyd 1992 using the complete body length from the anterior tip of the head to the posterior tip of the wing, excluding individuals that had truncated wings (<3% of their sample).

In 2013-2015 data, individuals with short wings made up approximately 48% of individuals from Key Largo and 4% of those from Lake Wales, and cannot be reasonably excluded from these samples. For individuals with truncated wings, this metric of body length is not a true indicator of the entire length of the body, as the abdomen extends beyond the tip of the closed wings regardless of an individual’s condition. I instead use thorax width to control for body size in 2013-2015 samples. Thorax width has a positive linear relationship with beak length in both populations (R^2^=.57) and with body length in long-winged individuals where both metrics were taken (R^2^=.77; Extended Data Fig. 6).

For comparisons including both 2013-2015 and 1988 data, I used a subset of individuals with both body length and thorax width measurements from 2013-2015 to create a ‘body length’ metric for all wing morphs using linear regression. This metric was generated by estimating the linear relationship between body length and thorax width (body length=2.796*thorax width +2.875) and using the thorax width of each individual to produce a body length for individuals without this measurement. Using this measure allowed the inclusion of overall size as a covariate in models comparing year and host effects on beak length. The top 3 models, which carried the majority of probability using AIC, all had weakly heteroscedastic residual errors. To compensate for this problem, a power transformation with λ=-0.1655463 was applied to beak length. This returned the same top 3 models, and did not alter the significance of any of the included factors or generate results different from those detected using pairwise t-tests.

Comparisons of all 8 populations measured in 2013-2015 were conducted using linear models or linear mixed models with all combinations of host, body size, and population as a random factor nested within host. The results of the top model, which carried >99% of the probability using AIC, are reported in the main text.

### Survival analyses

Historical survival means were extracted from Carroll et al 1998 Figure 4 using ImageJ. Using the sample sizes detailed in the same figure, the total number of survivors (1) and non-survivors (0) could be determined as a single possible number for each treatment, and so direct comparison to these data is possible. 95% confidence intervals in 2014 for individuals from *C. corindum* raised on *C. corindum* and *K. elegans,* respectively, are 0.17 & 0.22, while those from *K. elegans* are 0.18 and 0.22 when raised on the same two hosts. In 1988, the same 4 treatments have 95% confidence intervals of 0.27, 0.35, 0.32, and 0.24, respectively, using the Pearson-Klopper method. Confidence intervals for every population/natal host/year combination are plotted in Fig. 2 and Extended data Fig. 2.

For comparisons in only present day populations, all possible combinations of natal host, population host, individual population (random effect, nested within population host), family (random effect, nested within population), and interactions between fixed effects were considered in either generalized linear models (glm) or generalized linear mixed models (glmer in the package lme4) in the statistical program R using a binomial error distribution. All models were compared using AIC. For comparisons in only historical data, models containing natal host, population host, the interaction, and population as a random effect were compared using AIC. For clarity, results in the main text are reported from analyses conducted separately for each year.

For comparisons between years, all possible combinations of year, natal host, population host, individual population (random effect, nested within population host) and interactions between fixed effects were considered in either generalized linear models (glm) or generalized linear mixed models (glmer in the package lme4) in the statistical program R using a binomial error distribution. All models were compared using AIC. The results of the top 3 models for the between-year comparison are consistent with the results of within-year comparisons.

### Egg weight analyses

Historical egg weight data means and standard errors were extracted from Carroll et al 1998 Figure 2 using ImageJ. Using the sample sizes detailed in the same figure, data was simulated for statistical comparison to present day data. Data for each population was randomly sampled from a normal distribution with the mean and standard deviation of that population in 1988. This process was used to generate 1000 datasets with sample sizes equivalent to those in Carroll et al 1998. Present day egg weight data fails to reject normality using the Shapiro-Wilk normality test, making the normal distribution a reasonable choice for simulating historical data. Each simulated dataset was compared to the data collected in 2014 using two two-tailed t.tests: One looking at the effect of year on egg weight for bugs from *C.corindum* and one looking at the effect of year on egg weight for bugs from *K.elegans.* In all 1000 generated datasets, 2014 and 1988 measures were indistinguishable on *K.elegans* (t<1.76, df=48, p>0.05) and significantly different on *C. corindum* (t>5, df=18, p<.001).

For analyses of 2014 data alone, the generalized linear model of population host and a generalized linear mixed models (package lme4) containing individual and population were compared using AIC.

### Development time analyses

Historical development time data means and standard errors were extracted from Carroll et al 1997 Figure 3 using ImageJ. Using the sample sizes detailed in the same figure, data was simulated for statistical comparison to present day data. Data for each population and sex was randomly sampled from a normal distribution with the mean and standard deviation of that population and sex in 1991. This process was used to generate 1000 datasets with sample sizes equivalent to those in Carroll et al 1997. Present day development time data fails to reject normality using the Shapiro-Wilk normality test, making the normal distribution a reasonable choice for simulating historical data. To select the best linear model to describe these data, 10 sample datasets were each run through all possible linear models with combinations of the three main effects (host, year, and sex) and all possible interactions. These 19 models were then compared using AIC, and the model with the highest total weighted probability (calculated from AIC) across all ten iterations was selected. Each dataset was then compared to the data collected in 2014 using a this selected model (population host + year + sex + host*year + sex*year), along with two t.tests: One looking at the effect of year on egg weight for bugs from *C.corindum* and one looking at the effect of year on egg weight for bugs from *K.elegans.* The output of these models was then aggregated, and the summary of test statistics (t-values) and effect estimates was examined for each case. Mean effect estimates, t-values, and p-values are reported in the main text for pairwise t-tests. Predictors or interactions for which the effect size was in the same direction in >95% of cases (950 cases out of 1000) were considered to have a consistent, detectable effect. The results of these simulations are reported in Extended Data Table 1.

## Acknowledgements

I would like to thank L. Yang, C. Martin, T. Farkas, J. Berg, S. Carroll, and J. Rosenheim for their comments on earlier drafts of this manuscript. I would also like to thank S. Carroll for sharing his data and expertise, and A. Sharma for assistance with insect rearing. I thank the Florida State Parks Department for permitting the collection of bugs in Florida State Parks. Funding was provided by the University of California-Davis Department of Entomology, the Center for Population Biology, and a Henry A. Jastro Research Fellowship.

